# Characterization of HIV-1 particles co-purified with three extracellular vesicle subtypes from the Raji CD4 DCIR cell line, a hybrid model of CD4 T cells and dendritic cells

**DOI:** 10.1101/2025.03.27.645816

**Authors:** Julien Boucher, Alyssa Rousseau, Caroline Gilbert

## Abstract

**Background:** HIV-1 proteins and RNA are packaged into extracellular vesicles (EVs) through interaction with the multivesicular endosomes of the EV biogenesis machinery. This interaction also allows functional or abortive viruses to exit the cells as exosomes, microvesicles or apoptotic vesicles. HIV-1 viral particles and EVs share significant similarities in size, composition, and molecular cargo, making their separation challenging. Current HIV-1 purification methods neglect the effects of EVs on virus infectivity, which could influence experiment outcomes. Here, we co-characterized HIV-1 particles co-purified with exosomes, microvesicles and apoptotic vesicles to determine their impact on HIV-1 infection.

**Methods:** The HIV-infected Raji CD4 DCIR cells’ supernatants were harvested 2 and 8 days after infection. The 2-day supernatant was treated with proteinase K to discard viral protein and HIV-1 RNA associated with proteins outside the EVs. The supernatants were fractionated into three pellets by differential centrifugation. The 3K pellet contained the largest EVs, such as apoptotic vesicles. The 17K and 100K pellets were respectively associated with microvesicles and exosomes. EVs and viral particles were co-characterized for their host and viral contents and the pellets obtained after 8 days post-infection were tested for infectivity.

**Results:** Proteinase K treatment notably lowered HIV-1 RNA concentration in the EV pellet and did not affect viral p24 capsid protein concentration. The p24 protein was mostly found in the 17K pellet and HIV-1 RNA was the most abundant in the 100K pellet for both 2- and 8-day productions. Nevertheless, the 3K pellet had the highest infectivity when cells were infected with an equal quantity of virus (measured by p24) from each pellet.

**Conclusion:** Productively infected cells released functional viruses in the three EV subtypes, each exerting a distinct effect on virus infectivity. For future experiments, the presence of EVs in viral preparations should be taken into account, as they influence the progression of HIV-1 infection.

**Highlights:**

- HIV-1 RNA and p24 capsid protein are packaged within apoptotic vesicles, microvesicles and exosomes released by Raji CD4 DCIR cells.
- Infectious particles co-precipitated with the 3K pellet from Raji CD4 DCIR cells are more infectious than other viruses associated with other EV pellets.

## Introduction

HIV-1 infection targets the immune system to cause acquired immunodeficiency syndrome (AIDS) in people living with HIV-1 (PLWH) [1]. Fortunately, antiretroviral therapy (ART) prevents disease progression and AIDS development and successfully treated PLWH show an undetectable viral load in the peripheral blood [2]. However, latently infected cells establish reservoirs maintained despite long-term ART [3]. In ART-treated PLWH, viral replication remains active in lymph nodes [4]. Among the many factors influencing viral persistence, extracellular vesicles (EVs) are potent contributors to the ongoing pathogenesis [5].

EVs participate in cell-to-cell communication to influence many physiological processes, such as immunity [6]. Every immune cell releases a highly heterogeneous variety of EVs that can be categorized into three subclasses according to their biogenesis [6]. First, intraluminal vesicles are produced by membrane invagination in endosomes to form multivesicular bodies. The latter fuse to the cell membrane to release the EVs with a diameter of 30-150 nm, called exosomes, in the extracellular environment [7]. The second subclass includes vesicles, such as microvesicles (100-500 nm) and apoptotic vesicles (> 500nm), formed by cell membrane budding [8]. HIV-1 particles are similar to EVs in size, density and membrane composition [9]. Both particles acquire their membrane through the same mechanism, as HIV-1 hijacks the exosome biogenesis pathway to assemble viral particles [10]. Viral particles also assemble at the cell membrane, the microvesicle’s biogenesis pathway [11]. Finally, infected cells can go through apoptosis, releasing apoptotic vesicles that could contain viral material [12].

EVs integrate viral components to such an extent that they become virus-like particles (VLP) [9]. Transportation of viral proteins by EVs promotes HIV-1 pathogenesis. Nef-enriched EVs cause the apoptosis of CD4 T cells [13], increase the susceptibility of resting CD4 T cells to HIV-1 infection [14], and correlate with the systemic inflammation provoked by HIV-1 infection [15]. EVs can display the viral envelope protein gp120 on their surface [16]. EV-mediated gp120 enhances the infectivity of viral preparations [16]. Infected cells release Tat-enriched EVs, promoting neuronal death [17]. In addition, HIV-1 RNA is detectable in EVs of aviremic PLWH [18]. Quantifying HIV-1 RNA in large and small EVs of infected mice and PLWH showed a preferential enrichment in large EVs [19, 20]. Specific enrichment of HIV-1 RNA in large EVs is linked to immune activation and viral replication, suggesting a distinct contribution of EVs according to their subclass [20]. Finally, infectious viral particles have been detected in a wide range of EVs such as apoptotic bodies, microvesicles, exosomes and exomeres [21].

The quality and composition of HIV-1 viral preparations are crucial to testing antiviral drugs and deciphering HIV pathogenesis. HIV-1 exploits different pathways to produce viral particles, which introduce heterogeneity in their composition. The natural integration of specific host proteins on viral particles alters viral infectivity [22–24], potentially influencing the outcome of experiments to understand the infection pathogenesis and drug efficacy studies. Given this complexity, the proportion of infectious versus non-infectious viral particles within different EV subtypes and their relative contribution to HIV-1 pathogenesis remains unclear. This gap in knowledge underscores the need for optimized purification methods to ensure that laboratory-prepared HIV-1 particles accurately reflect the biologically relevant virus required for robust in vitro and in vivo experiments.

The traditional in-laboratory HIV-1 stock production method is human embryonic kidney (HEK293T) cell transfection with a plasmid containing the HIV-1 genome [25]. Due to a divergent production pathway, transfected cells and infected cells produce viral preparations with different characteristics. For example, HIV-pulsed PBMCs release miR-155-rich EVs that promote HIV-1 replication [26]. HEK293T cells release do not express immune factors such as miR-155, HLA-DR and ICAM-1 [26–28]. Thus, virus stock by cell infection could be a better option to recapitulate infection events in experimental settings.

In this study, we sought to produce a more physiologically relevant HIV-1 stock by infecting a Raji CD4 DCIR cell line. Raji CD4 cells are B cells, transformed to express CD4, rendering them susceptible to HIV-1 infection [29]. Raji CD4 cells were further modified to express the dendritic cell immunoreceptor (DCIR), an HIV-1-binding lectin expressed by dendritic cells [30, 31]. Thus, Raji CD4 DCIR generates a hybrid model of lymphocytes and dendritic cells to study HIV-1 infection by X4-tropic virus. We characterized the heterogeneity of viral particles and EVs produced by infected cells 2 and 8 days post-infection. The supernatant containing EVs and viruses, or EVs only from non-infected cells, was fractionated into three pellets: 3K, 17K and 100K, which should correspond to apoptotic vesicles, microvesicles and exosomes [32, 33]. Characterizing viral components in each fraction showed that the concentration of viral protein and viral RNA was higher in the 17K and 100K pellets. The most infectious viruses produced by infected Raji CD4 DCIR were found in the 3K pellet and the least infectious in the 100K pellet.

## Methods

### HIV-1 stock preparation by HEK293T transfection

NL4-3 (X4 tropic) (AIDS Repository Reagent Program) virus was produced in HEK293T by transient calcium/phosphate transfection, as described previously [34]. To produce HIV-1 particles, 2 x 10^6^ HEK293T cells were transfected with 20 µg of a pNL4.3 plasmid. Then, cells were washed 16 hours after transfection and kept in culture in DMEM supplemented with 2% decomplemented and ultracentrifuged FBS for 48 hours. The cell-free supernatant was filtered at 0.20 µm and then centrifuged at 100,000 × *g* for 45 minutes (100K pellet) in an Optima L-90K Beckman Coulter centrifuge with a 70 Ti rotor to pellet virus and EVs.

### Infection of Raji CD4 DCIR cells

Two different lots of Raji CD4 DCIR were incubated with 100 ng of NL4.3 p24 per 10^6^ cells for 2 hours. Control cells were incubated with an equivalent volume of PBS. Cells were washed three times with PBS to remove the free virus and maintained in culture in X-VIVO medium (Lonza) to obtain a concentration of 10^6^ cells/mL. The cell culture with X-VIVO is FBS-free, thus avoiding the bovine EV contamination of our EV/HIV-1 productions. Two days post-infection, the whole cell culture supernatant was harvested and replaced with fresh X-VIVO medium. The two-day supernatant was treated overnight with AT-2 (2, 2′-Dithiodipyridine) to inactivate HIV-1 particles before EV/HIV-1 purification. Eight days post-infection, the whole supernatant was harvested for EVs and virus co-purification. The experiment was repeated twice with two different lots of Raji CD4 DCIR to obtain four EV/HIV-1 production replicates.

### Raji CD4 DCIR EV/HIV-1 purification

Before the purification protocol, a portion of the two-day supernatant was treated with proteinase K at a final concentration of 1.25 mg/mL for 10 minutes at 37 °C. Proteinase K treatment eliminates extracellular lipoproteins and RNA-binding proteins on the EVs to quantify only the intravesicular HIV-1 RNA and viral and host proteins. Then, EV/HIV-1 were co-purified from the two-day and eight-day supernatants following the same procedures. The supernatants were centrifuged at 3,000 x *g* for 15 minutes (3K pellet), then at 17,000 x *g* for 30 minutes (17K pellet) and finally at 100,000 x *g* for 60 minutes (100K pellet) (Figure 1). All pellets were resuspended in 0.20µm-filtered PBS 1X in 1/10^th^ of the original volume. EVs were immediately stored at -80°C until further analysis.

**Figure 1.**
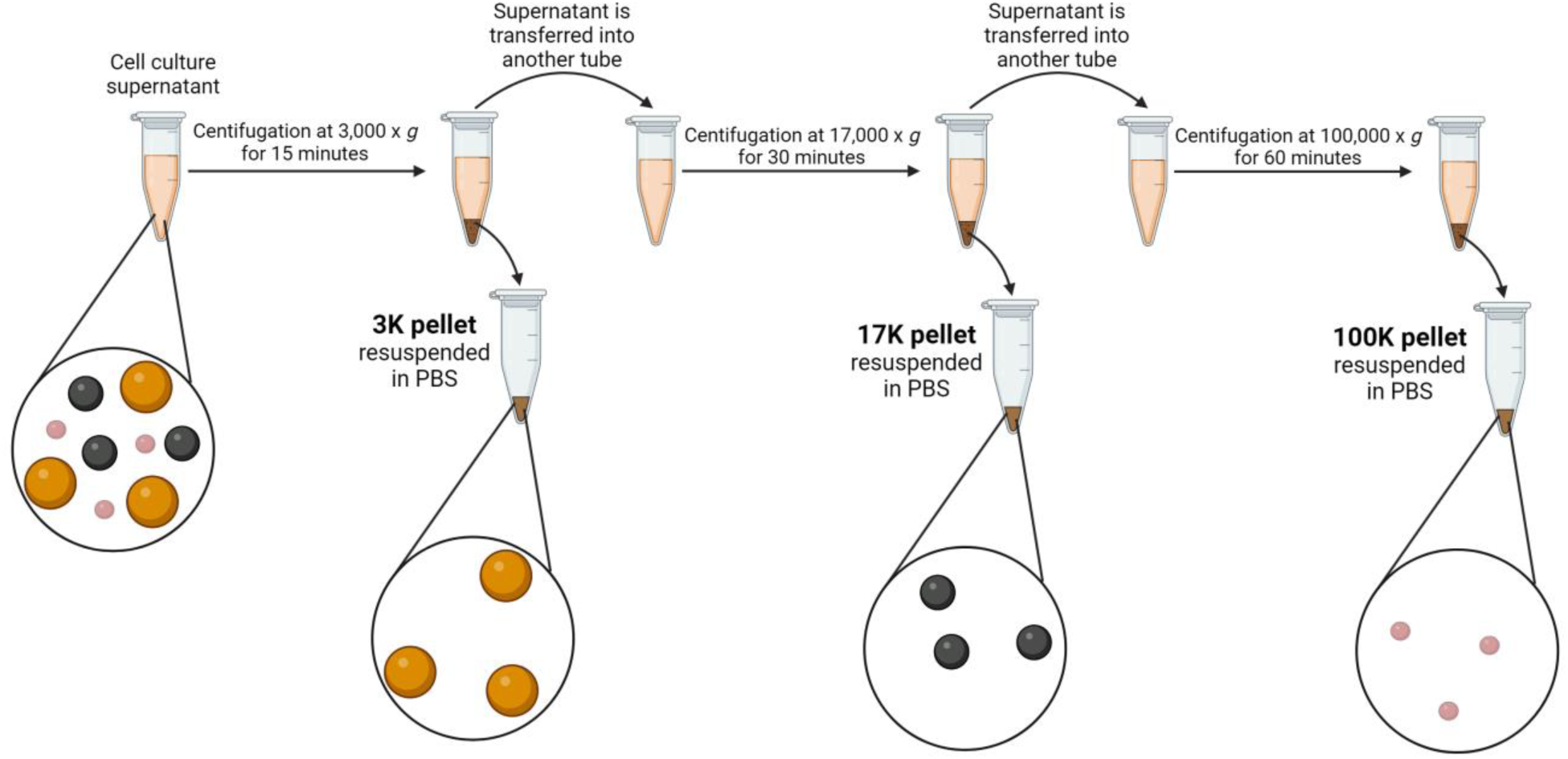
EVs and viruses purification protocol. Schematic representation of the differential centrifugation steps to obtain the 3K, 17K and 100K EV and virus pellets.

### Hydrodynamic size measurement

EVs size was analyzed by dynamic light scattering (DLS) with a Zetasizer Nano ZS (Malvern Instruments) as described before [26]. Hydrodynamic diameter measurements were done in duplicate at room temperature. The derived count rate provided an estimation of EV concentration. This measure indicates the number of photons scattered by nanoparticles (shown in kilo count per second; kcps).

### EVs quantification by flow cytometry

Absolute EV quantification by flow cytometry was performed as previously described [35]. EVs were stained with lipophilic carbocyanine DiD dye and CellTrace^TM^ CSFE (ThermoFisher Scientific), both at a final concentration of 5 µM. DiD+ and CSFE+ events were considered EVs. To determine the EV concentration in our samples, a fixed quantity of 2µm Cy5-labelled silica beads (Nanocs), measured before every experiment by using a Cellometer, was added to every sample before cytometer acquisition. The number of acquired beads during the cytometer experiments allowed us to calculate the analyzed volume. Then, we divided the absolute number of EVs acquired by the cytometer by the analyzed volume to determine the EV concentration. The gating strategy was previously described [20].

### Transmission electron microscopy

Briefly, 100 µL of purified EVs were diluted to 1 mL with 2.5% glutaraldehyde plus sodium cacodylate 0.1 M pH7.4. EVs were incubated at 4°C for 1 hour for fixation. Then, EVs were centrifuged with the same parameters described in the EVs purification section. Pellets were suspended in 25 µL sodium cacodylate 0.1 M pH 7.4 and kept at 4°C until grid preparation. A mix of 2 µL of the sample and 2 µL of 2% uranyl acetate was pipetted onto a copper grid with carbon support film. The grids were stained for 40 seconds. Excess liquid was removed by blotting and incubation at 37°C O/N. Images were acquired at 80 kV using a JEOL 1230 (JEOL, Tokyo, Japan) transmission electron microscope with a bottom-mounted CCD camera.

### Anti-p24 ELISA

The viruses were titrated for all stock preparations and experiments using an in-house ELISA for the viral protein p24 [36].

### Western Blot

The resuspended EV/HIV-1 preparations (50 µL) were added to 12.5 µL of 5X Laemmli sample buffer and heated at 100 °C for 7 minutes. A total of 15 µL was loaded on a 10% sodium dodecyl sulfate-polyacrylamide gel to perform electrophoresis. The separated proteins were transferred onto an Immobilion® PVDF membrane. The PVDF membrane was blocked in 5% powdered milk dissolved in PBS-T (PBS with 0.1% Tween-20). Primary antibody incubation was carried out overnight at 4°C with continuous agitation. Anti–ICAM-1 (G-5, diluted at 1/200), anti–HSP70 (W27, diluted at 1/200), anti–TSG101 (C-2, diluted at 1/100) and anti–DAP-3 (C10, diluted at 1/100) were purchased from Santa Cruz Biotechnology Inc. Anti–HLA-DR (DA6.147, diluted at 1/200) and anti–p24 (183-H12-5C, diluted at 1/200) were produced in the laboratory from hybridoma cell lines. The 183-H12-5C and DA6.147 hybridomas were obtained through the AIDS Repository Reagent program and ATCC, respectively. The antibodies were purified using a HiTrap Protein G column (GE Healthcare). The membranes were washed three times in PBS-T and incubated with a 1/20,000 dilution of horseradish peroxidase (HRP)-conjugated anti-mouse secondary antibody purchased from Jackson ImmunoResearch. The signal was revealed on a HyBlot® autoradiography film (Thomas Scientific) with the Lumina Forte HRP substrate.

### Virus infectivity assay with a luciferase-based reporter cell line

To titrate virus stock infectivity, we used the TZM-bl indicator cell line, which carries a stably integrated luciferase reporter gene under the control of the HIV-1 regulatory element (LTR).

Following cell contact with the produced virus, we measured the replication level of a competent infectious virus using luciferase fluorescence detection [20]. A fixed volume of virus preparation or virus quantity (measured by p24) was added to a 96-well plate. Then, a serial dilution with a two-fold factor was performed to obtain 12 dilutions in triplicate. Cells were added (10,000 cells / well) and incubated for 48 hours at 37 °C, 5 % CO_2_. Then, cells were lysed with luciferase lysis buffer (25 mM Tris phosphate, pH 7.8, 2 mM dithiothreitol, 1 % Triton X-100, and 10 % glycerol) and frozen at -20°C. Twenty-four hours after lysis, an aliquot of cell extract was mixed with luciferin buffer (20 mM tricine, 1.07 mM magnesium carbonate hydroxide pentahydrate, 2.67 mM magnesium sulphate, 100 µM EDTA, 220 µM Coenzyme A, 4.70 µM D-Luciferin potassium salt, 530 µM ATP, and 33.3mM dithiothreitol) and luciferase reaction was measured using a Varioskan luminometer (ThermoFisher). The TCID_50_ (50% Tissue culture infectious dose) was finally calculated with the Spearman-Karber method [37].

### Infection of PBMCs

The CHU de Québec research ethics committee received prior approval for this study. All participants provided written consent. Peripheral blood mononuclear cells (PBMCs) were isolated from healthy donors on a lymphocyte separation medium [26]. PBMCs were washed and resuspended in HBSS (Hank’s balanced salt solution). Cell viability was determined by trypan blue coloration. PBMCs were distributed in 12-well plates at 10^6^/mL in X-VIVO medium. Cells were incubated with HIV-1 from the 3K, 17K or 100K pellet (20 ng of p24 per million cells) for 2 hours, then washed three times, resuspended in complete RPMI, and cultured for 4 days at 37°C under 5% CO_2_. Cells were resuspended in Trizol reagent for RNA extraction and in lysis buffer (Tris-HCl pH 8.3 0.1M, Tween 20 0.1%, Triton X-100 0.1% and proteinase K 400 µg/mL) for DNA extraction.

### Iodixanol velocity gradient

The 3K, 17K and 100K pellets were further purified on an iodixanol (OptiPrep) velocity gradient of 11 fractions, ranging from 6.0 to 18.0% iodixanol with 1.2% increments [38]. The gradient was centrifuged at 180,000 x *g* for 50 minutes with slow acceleration and no brakes in a StepSaver 65V13 vertical rotor. The fractions (900 µL) were immediately harvested after centrifugation. For RNA extraction, 100 µL of each fraction was diluted in 300 µL of Trizol LS (ThermoFisher Scientific) and 100 µL of each fraction was diluted in 300 µL of DNA lysis buffer (Tris-HCl pH 8.3, 0.1M, Tween 20 0.1%, Triton X-100 0.1% and proteinase K 400 µg/mL) for DNA extraction. The remainder was used for capsid p24 quantification with an in-house ELISA and total protein quantification with a micro BCA™ protein assay kit (ThermoFisher Scientific).

### RNA extraction

RNA was extracted from cells (resuspended in Trizol®) and from their EVs (in suspension diluted in 3 volumes of Trizol® LS) using the phenol/chloroform method [26]. Its concentration was measured using a BioDrop™ spectrophotometer. Its quality was considered suitable when the 260/280 nm absorbance ratio was between 1.5 and 2. Total RNA was treated with DNase I (0.2 U/µL) in 10 µL of DNase buffer (10 mM Tris, pH 7.5; 2.5 mM MgCl2; 0.5 mM CaCl2) at 37°C for 20 minutes. Then, DNase was inactivated at 65°C for 10 minutes after adding 1 µL of EDTA 50 mM.

### DNA extraction

Genomic DNA was extracted from PBMCs by the phenol/chloroform method. The lysate was treated with proteinase K (1 mg/mL) for 5 minutes at 55°C. Phenol/chloroform/isoamyl alcohol was added, followed by centrifugation at 12,000 × *g* for 3 minutes. Chloroform was added to the aqueous phase, followed by centrifugation at 12,000 × *g* for 3 minutes. DNA was precipitated from the second aqueous phase at -20°C by adding absolute ethanol, 0.5M NaCl, and GlycoBlue Coprecipitant (30 µg/mL final concentration), with centrifugation at 12,000 × *g* for 15 minutes. DNA concentration was measured with a BioDrop™ spectrophotometer. DNA quality was considered suitable for subsequent analysis when the 260/280 nm absorbance ratio was between 1.8 and 2. DNA extract was diluted 10-fold for downstream analysis.

### Viral load quantification

The total RNA extracted by the phenol/chloroform method was resuspended in 15 µL of Tris/EDTA buffer. RT-PCR was performed on 5 µL of RNA with Superscript IV reverse transcriptase according to the manufacturer’s instructions on a GeneAmp® PCR System 9700 (Applied Biosystems). The primer set targets the HIV-1 LTR region shown in Table S2 [39]. Reactions were performed with 1 µL of 5-fold diluted cDNA on a CFX384 Touch Real-Time PCR Detection System (Bio-Rad). The precision of our HIV RNA quantification method was validated with a quantification standard (NIH AIDS Reagent Program). For HIV-1 RNA quantification in plasma EVs, cDNA was pre-amplified to increase qPCR sensitivity, as previously established [20].

### Quantification of HIV DNA

The pre-amplification and qPCR reactions for total and integrated HIV genome copies and CD3 gene quantification were performed as described previously with the same primer sets [40], but using DNA purified by the method described above. For all qPCR reactions, TaqMan™ Fast Advanced Master Mix (Invitrogen) was used. All pre-amplification reactions were performed on a GeneAmp® PCR System 9700 (Applied Biosystems), and all qPCR reactions were performed on a CFX384 Touch Real-Time PCR Detection System (Bio-Rad).

### Quantification of mitochondrial DNA and KRAS DNA

Mitochondrial DNA was quantified by qPCR as described before [41]. Briefly, 1 µL of DNA, normalized at 2 ng/µL, was mixed with 5 µL QuantiTech SyBr Green, 2 µL of PCR grade water and 1 µL of each primer. The primers were obtained from Integrated DNA Technologies: forward: 5’-ACGCCTGAGCCCTATCTATTA-3’; reverse: 5’-GTTGACCTGTTAGGGTGAGAAG-3′.

KRAS DNA was quantified as a control of total gDNA. The primers were obtained from Integrated DNA Technologies: forward: 5’-CCTTGGGTTTCAAGTTATATG-3’; reverse: 5’-CCCTGACATACTCCCAAGGA-3′.

### Statistical analysis

Statistical analyses were carried out using GraphPad Prism software version 10.1.1 with p-values below 0.05 considered statistically significant. Two-way ANOVA was performed to compare the four EV/HIV-1 preparations in multiple conditions. Flow cytometry data were analyzed using FlowJo software version 10.6.1.

## Results

### HIV-1 RNA and p24 capsid protein in EVs released after a 2-day infection are partially protected from proteinase K treatment

HIV-1-infected cells release EVs bearing viral components before mature viral particles [42]. Here, the distribution of viral components in three EV subtypes was assessed. The culture supernatants from infected Raji CD4 DCIR or non-infected Raji CD4 DCIR (Mock) were harvested after two days and treated with proteinase K to validate that measured viral components were in the lumen of EVs. Culture supernatant EVs were fractionated by sequential centrifugation to obtain 3K, 17K and 100K pellets (Figure 1). EVs’ hydrodynamic size was measured to determine the size and heterogeneity of each preparation of EVs and HIV-1 (Figure 2A). DLS analysis showed that proteinase K pre-treatment modestly reduced the size and heterogeneity of EVs, except for the mock 3K and 17 K EVs, where proteinase K pre-treatment had a greater effect (Figure S1A-B). Relative EV concentrations were estimated with the derived count rate, and proteinase K decreased the count for the 3K and 17K pellets (Figure 2B). HIV-1 infection did not affect EV release by Raji CD4 DCIR cells (Figure 2B). Flow cytometry was used to obtain an absolute concentration of EVs. The impact of the proteinase K on EVs concentration decrease was preferentially observed in the 17K and 100K pellets (Figure 2C). HIV-1 infection was associated with an EV concentration increase in the 17K pellet and a decrease in the 100K pellet (Figure 2C).

**Figure 2.**
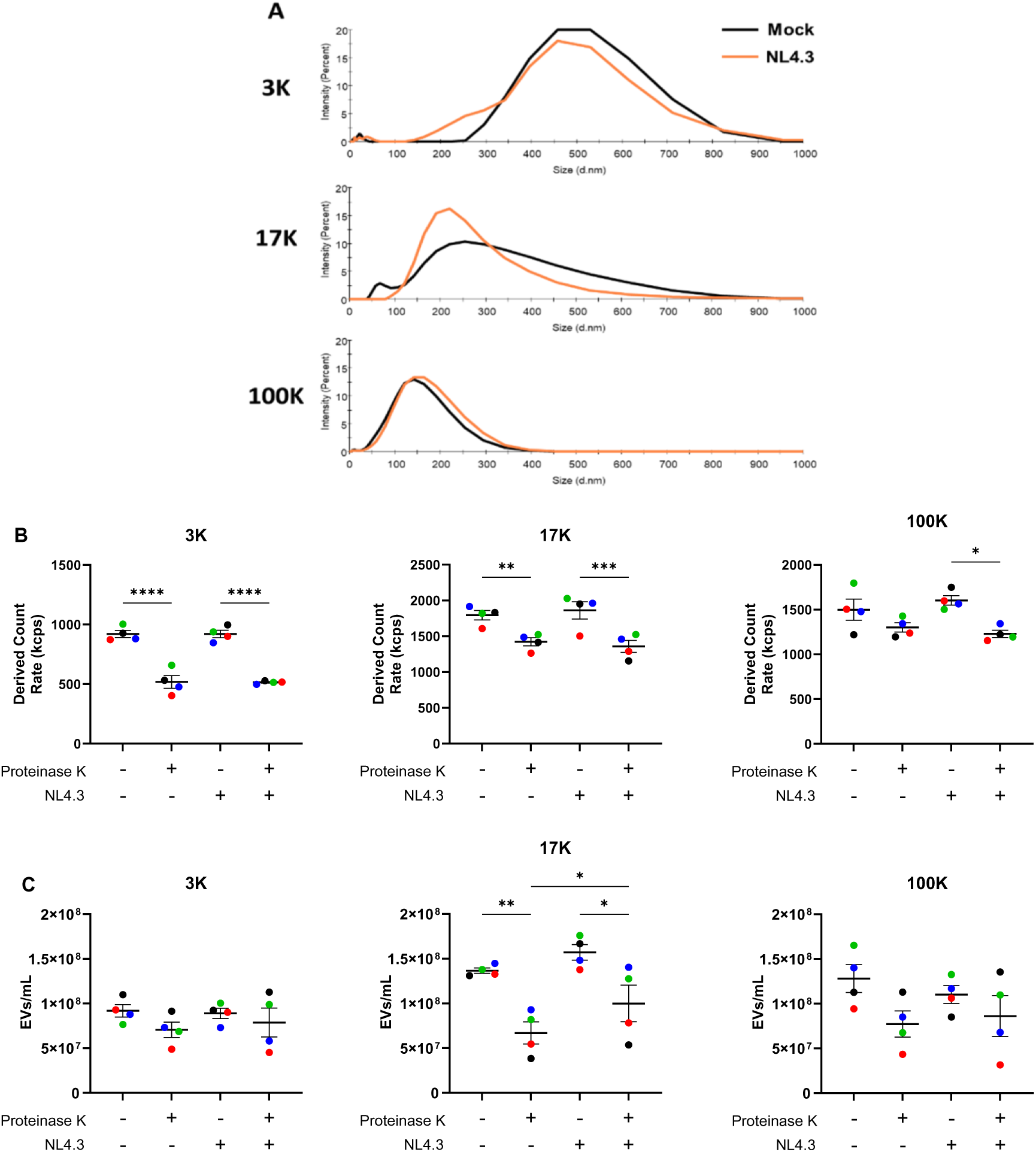
HIV-1 infection does not change the size and abundance of Raji CD4 DCIR-derived EVs 60 hours post-infection. Hydrodynamic size **(A)** and derived count rate **(B)** were measured by DLS. Absolute EV concentration was obtained by flow cytometry **(C)**. Data represent the mean of EVs from four different lots of Raji CD4 DCIR cells (n = 4). Statistical analysis was carried out by two-way ANOVA followed by Tukey’s multiple comparison post-test (* p < 0.05; ** p < 0.01; *** p < 0.001; **** p < 0.0001).

Proteinase K treatment effectiveness was determined by Western blot analysis of ICAM-1 and HSP70. ICAM-1 localization at the membrane makes it susceptible to proteinase K digestion [43]. HSP70 is found in the lumen of EVs and, therefore, should be protected from proteinase K treatment [44]. ICAM-1 was lowered in proteinase K-treated EVs, while HSP70 presence was maintained (Figure S2).

We quantified viral RNA and capsid protein p24 in the EV subtypes, purified from supernatant treated or not with proteinase K. HIV-1 RNA was the most abundant in the 100K EVs, but considerable concentrations were measured in the 3K and 17K pellets (Figure 3A). Proteinase K reduced viral RNA concentrations in all pellets, with the strongest effect on the 100K pellet. Moreover, proteinase K treatment changed the distribution of HIV-1 RNA in the three pellets. The 17K pellet contained slightly more viral RNA than the 100K pellet (Figure 3A). The p24 protein distribution in the pellets was vastly different than viral RNA. The 17K contained significantly more p24 than the 3K and the 100K pellets. Proteinase K did not affect p24 quantification (Figure 3B). HIV-1 RNA and p24 capsid protein concentrations in EVs were expected to correlate since both are viral components. For non-treated EVs, no correlation was found, while proteinase K treatment resulted in a weak, but not significant correlation (Figure 3C). These results suggest that a considerable amount of HIV-1 RNA in EVs is not associated with p24 production. Next, we estimated the viral particle concentration using the results of HIV-1 RNA and p24 quantification assays. There are two RNA copies per HIV-1 particle [45]. Thus, viral particle concentration can be estimated by dividing HIV-1 RNA concentration by a factor of two. Similarly, there are approximately 3,000 p24 molecules per virus, which allows the calculation of 8.36 * 10^6^ particles per ng of p24 [45]. Similar proportion of viral particles were calculated with both measurements in non-treated supernatants, except the 17K pellet (Figure 3D). This calculation also showed that the p24 measurement estimates more viral particles in the proteinase K pre-treated pellets (Figure 3E). These results suggest the presence of extravesicular HIV-1 and genomic RNA-free viral capsids.

**Figure 3.**
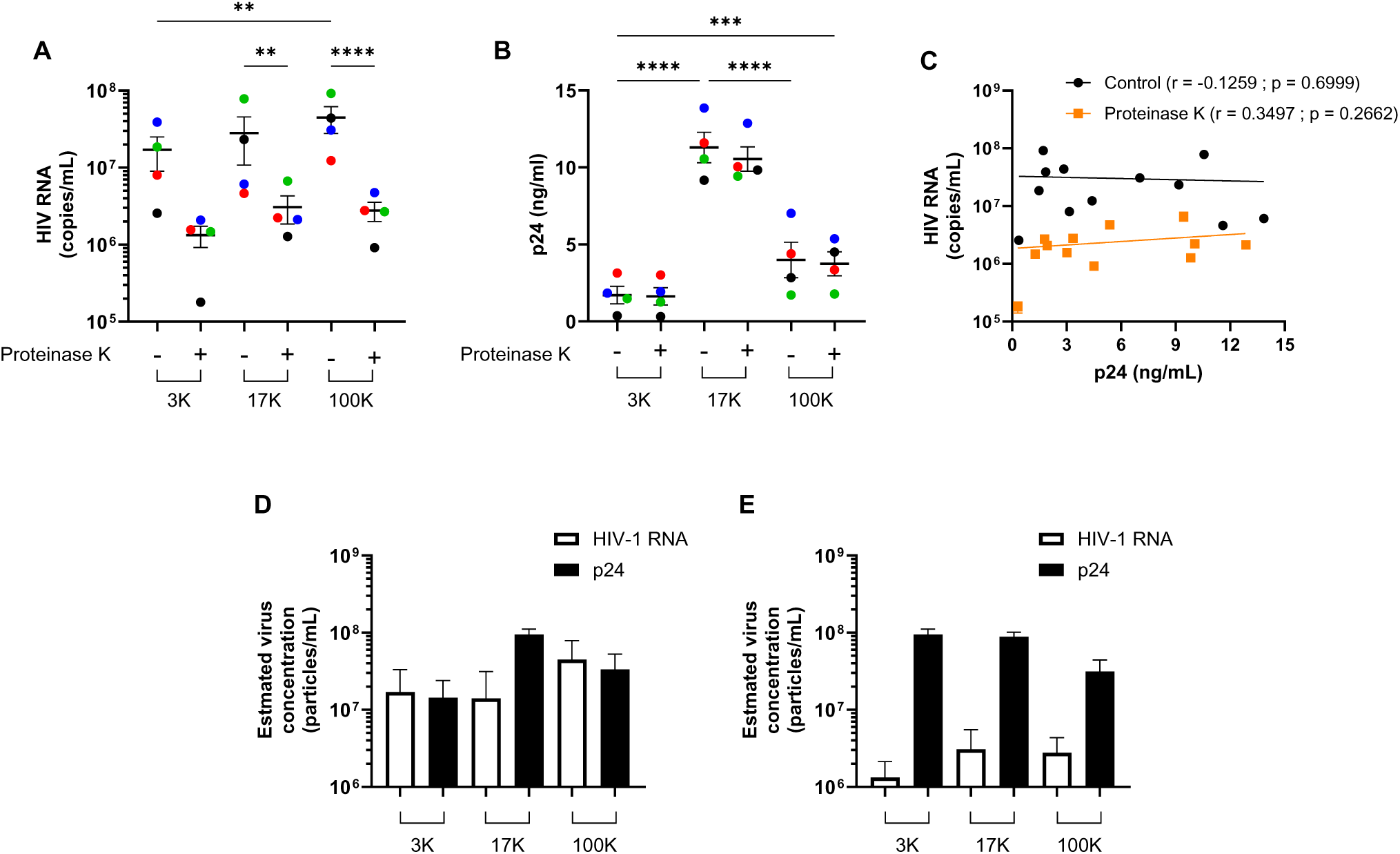
Distribution of viral components in EVs during early infection. HIV-1 RNA was quantified by RT-qPCR **(A)**. Capsid protein p24 was quantified by ELISA **(B)**. Spearman correlations between HIV-1 RNA and p24 concentrations in the non-treated and proteinase K-treated EVs **(C)**. Viral particles concentration estimated with the HIV-RNA and the p24 concentration in non-treated supernatant **(D)** and in the proteinase K-treated supernatant **(E)**. Data represent the mean of EVs from four different lots of Raji CD4 DCIR cells (n = 4). Two-way ANOVA and Tukey’s multiple comparisons test were used for statistical analysis (**p<0.01, ***p<0.001, ****p<0.0001).

### Infectivity of virus associated with EV subtypes

We established that viral proteins and RNA are associated abundantly with all EV subtypes. Thus, we maintained infected Raji CD4 DCIR cells in culture for 8 days to produce infectious viruses to determine infectivity distribution within the three EV subtypes. Purified EVs and viruses were analyzed by DLS to measure their hydrodynamic size. As expected, DLS measurement showed that the 3K pellet remained the largest and the 100K was the smallest (Figure 4A), which was also confirmed by transmission electron microscopy (Figures 4B and S3). EV quantification by flow cytometry showed that the most EVs were precipitated in the 100K and the least in the 3K pellet (Figure 4C). As expected, mitochondrial DNA in EVs was at a higher concentration in larger EVs (Figure 4D). We also measured KRAS (Kirsten rat sarcoma viral oncogene homologue) DNA abundance in EVs as a control of gDNA distribution. We measured that the 100K pellet contained the most KRAS DNA (Figure 4E). Finally, HIV-1 RNA was the most abundant in the 100K pellet (Figure 4F), while p24 was the most abundant in the 17K pellet (Figure 4G). Again, HIV-1 RNA and capsid p24 protein concentrations in the EV/HIV-1 preparations did not correlate (Figure 4H). The virus particle concentration in the 8-day production was estimated with the same formula described above. The estimated viral particle concentration by HIV-1 RNA was lower than the estimation with the p24 viral protein (Figure 4I). This data suggests a surplus of p24 production compared with HIV-1 RNA availability to build complete viral particles.

**Figure 4.**
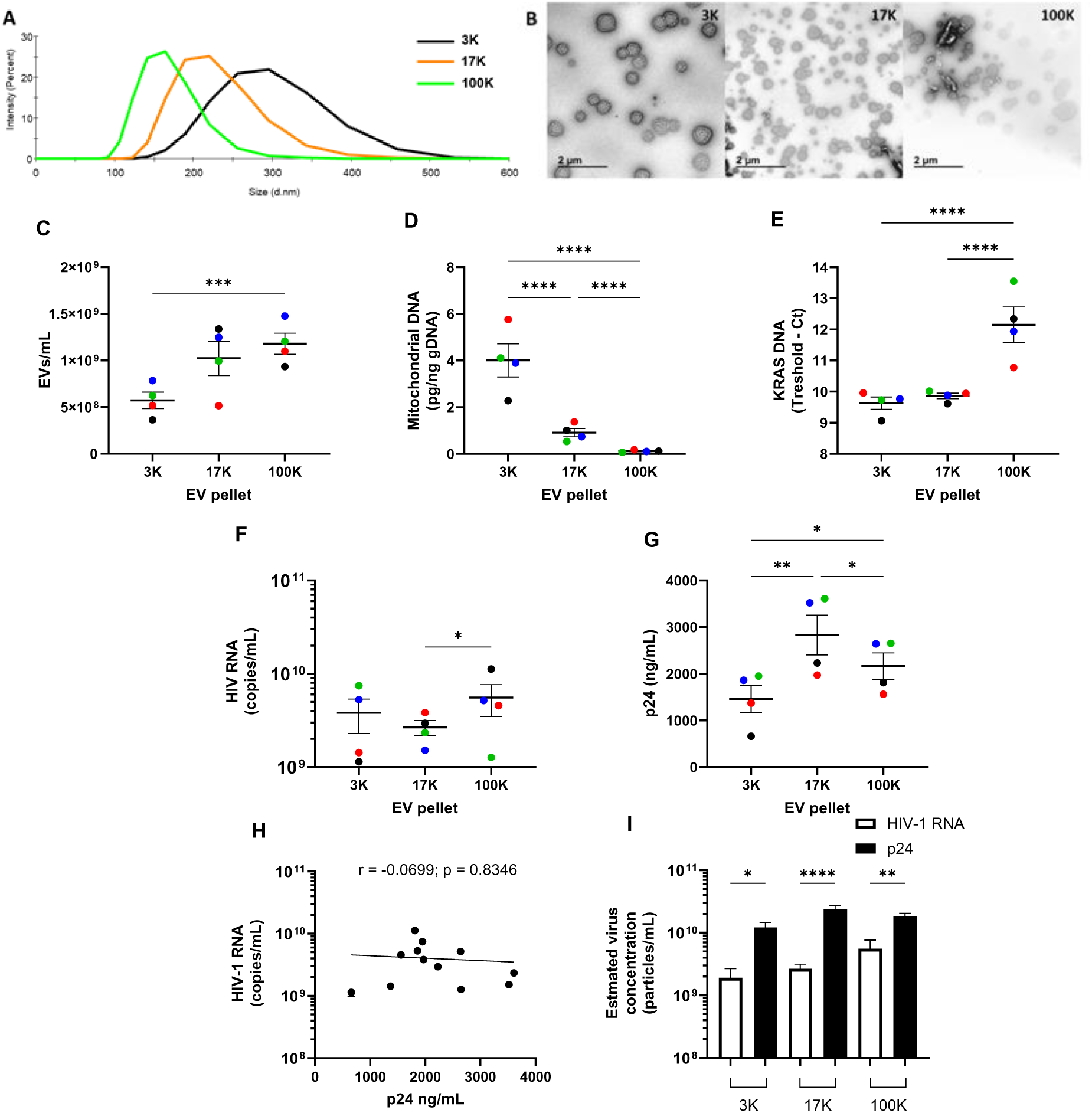
Characterization and distribution of viral components in EV/HIV-1 pellets. Hydrodynamic size measurement of the three purified EV/HIV-1 fractions **(A)**. Transmission electron microscopy at a 3,000X magnification of pelleted EVs **(B)**. Absolute nanoparticle concentration was obtained by flow cytometry **(C)**. Mitochondrial DNA quantification in each fraction by qPCR. Results are presented as pg of mitochondrial DNA per ng of gDNA **(D)**. KRAS DNA expression measured by qPCR. The threshold is defined as 38, which is the lowest Ct value to be considered as a valid result **(E)**. Viral load quantification in each fraction by RT-qPCR **(F)**. Capsid protein p24 quantification in each fraction by ELISA **(G)**. Spearman correlations between HIV-1 RNA and p24 concentrations **(H)**. Viral particles concentration estimated with the HIV-RNA and the p24 concentration **(I)**. Data represent the mean of four different lots of Raji CD4 DCIR cells (n = 4). Two-way ANOVA with Tukey’s multiple comparison test was carried out for statistical analysis (* p < 0.05; ** p < 0.01; **** p < 0.0001).

EV and virus pellets were further characterized by assessing their protein content (Figure 5). HSP70 is a ubiquitous protein detectable in exosomes and microvesicles [46]. Here, HSP70 was mostly associated with the 17K pellet but was also present in the 3K and 100K pellets. Being a membrane protein, ICAM-1 is displayed by microvesicles [43]. ICAM-1 was strongly detected in the 17K pellet. DAP-3 is a mitochondrial protein involved in apoptosis and was expected to be detected in apoptotic vesicles, thus, the 3K pellet [47]. The strongest signal was detected in the 3K pellet, with a weak signal from the 17K pellet (Figure 5). Since the Raji cells are B cells, they express HLA-DR. As previously observed in dendritic cells, the highest HLA-DR detection was anticipated in exosomes [33]. The 100K pellet displayed the highest abundance of HLA-DR.

**Figure 5.**
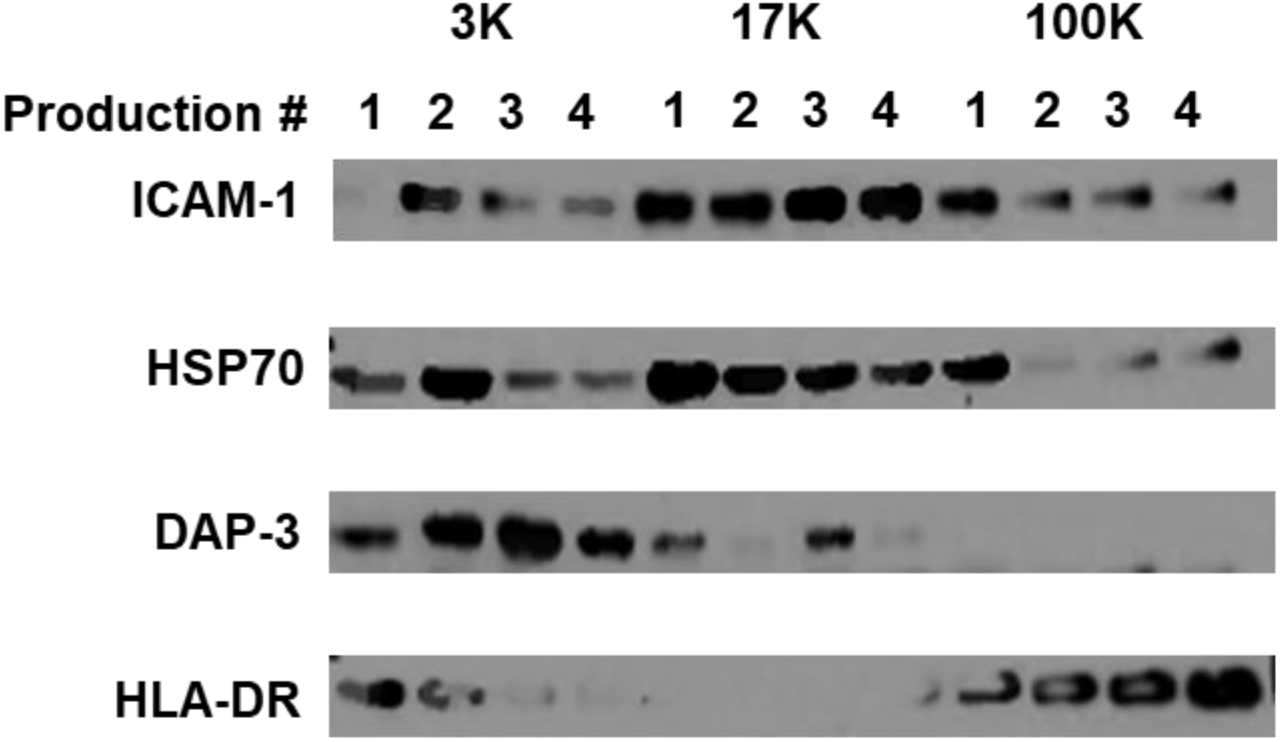
EV protein analysis by Western blot. Characterization of EV-associated proteins in the 3K, 17K and 100K EVs and virus pellet from infected Raji CD4 DCIR cells from four independent experiments.

The infectivity of the virus in the 3K, 17K, and 100K fractions was initially tested on a TZM-bl reporter cell line. An equivalent amount of virus (20 ng of p24) was used to infect cells in every condition. Unexpectedly, viruses from the 3K pellet were significantly more infectious, while viruses from the 100K pellet were the least infectious (Figure 6A and S4). To confirm these results, PBMCs were infected with the same viruses at a concentration of 20 ng of p24 per million cells. Four days after infection, intracellular HIV-1 RNA was quantified, and the 3K pellet was associated with the highest viral RNA production (Figure 6B). To validate that viruses from the 3K pellet were more infectious, integrated (Figure 6C) and total (Figure 6D) intracellular viral DNA was quantified. Both HIV-1 DNA forms were at the highest concentration when PBMCs were infected with the 3K pellet. Viruses from the 100K pellet were the least infectious. The hydrodynamic size measurement (Figure 4A), the transmission electron microscopy observations (Figures 4B and S3) and the flow cytometry analysis (Figure S5) confirmed the absence of cell-sized particles. These results suggest that HIV-1 particles associated with the 3K pellet are more infectious than those co-purified with the 17K and 100K pellets.

**Figure 6.**
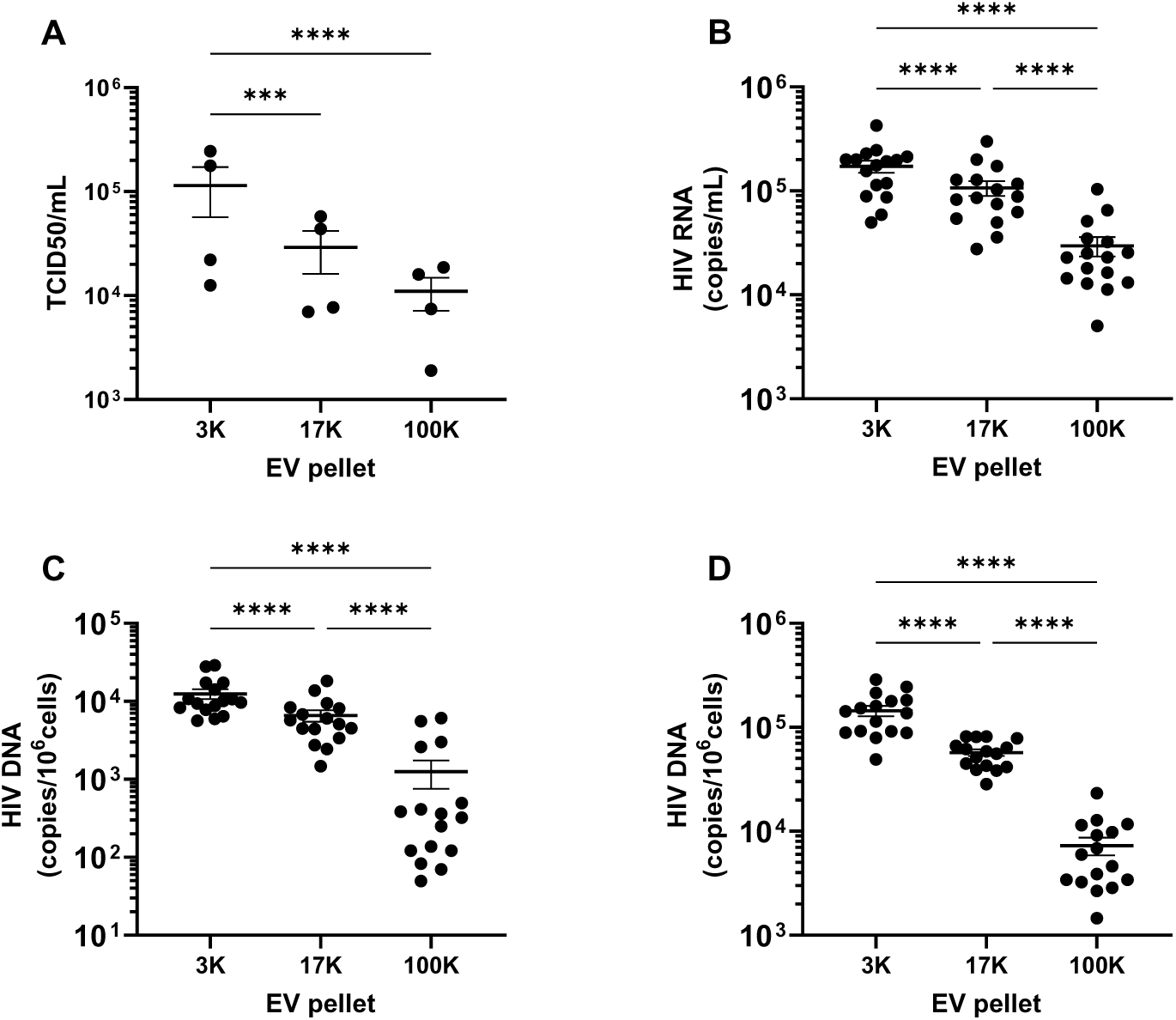
Infectivity of HIV-1 particles associated with three EV fractions. Infectivity of HIV-1 particles in the 3K, 17K and 100K pellets was assessed with a TZM-bl indicator cell line. An equal amount of virus (equivalent to 20 ng of p24) was used to infect the cells **(A)**. Freshly purified PBMCs from four non-infected donors (n = 4) were infected with an equal amount of virus (20 ng of p24/million cells) from the 3K, 17K or 100K pellets of four distinct viral preparations. Four days post-infection, intracellular HIV-1 RNA was quantified by RT-PCR **(B)**. Integrated HIV-1 DNA in the PBMC’s genome was quantified by a nested qPCR **(C)**. Total intracellular HIV-1 DNA was quantified by qPCR **(D)**. Two-way ANOVA with Tukey’s multiple comparison test was performed for statistical analysis (*** p < 0.001; **** p < 0.0001).

HEK293T cell transfection with a plasmid coding for the whole HIV-1 genome is a common method used for viral stock production. Transfected HEK293T released a negligible quantity of viruses associated with the 3K pellet (Figure S6), whereas the 3K pellet from infected Raji CD4 DCIR contained the most infectious particles. This highlights the divergence of viruses produced by transfected or infected cells. The virus particle concentration in the HEK293T production was also estimated using the formula described above. Again, these calculations showed a p24 estimation 5 to 15 times higher than the estimation with the HIV-1 RNA measurement (Figure S6D).

### Ultra-purification of EVs and viruses with the iodixanol velocity gradient

The velocity iodixanol gradient is an established procedure to separate HIV-1 particles and EVs [38]. Raji CD4 DCIR-derived EVs and viruses were processed on iodixanol velocity gradients to separate viruses and EVs harboring viral components. The gradient consists of 11 fractions ranging from 6.0 to 18.0% iodixanol with 1.2% increments. Infectious viruses are mostly expected in the 14.4 to 18.0% fractions, while EVs are found in the 9.6 to 12.0% fractions [38]. Total RNA and DNA were not preferentially enriched in specific fractions of the gradients (Figure S7A). Total proteins were slightly more abundant in the denser fractions (Figure S7B). Mitochondrial DNA was mostly found in the 18.0% fraction of the 3K and 17K pellets (Figure S8A). MiR-155 profile was constant in the three gradients, where a strong proportion was in the EVs fractions (7.2 to 10.8%) and the virus fractions (15.8 to 18.0%) (Figures S8B).The viral content was analyzed for each fraction to find enrichment in fractions 15.8 to 18.0% as previously observed [38]. Interestingly, the 3K pellet had a higher proportion of p24 protein to viral RNA (Figure 7A), which was not observed in the 17K (Figure 7B) and 100K pellet (Figure 7C). Virus and EVs separation on the iodixanol velocity gradient revealed that HIV-RNA and p24 protein are mostly detected in the virus fractions (14.4 to 18.0%). Finally, the virus particle concentration was estimated in the 14.4 to 18.0 % fractions of iodixanol velocity gradient. This estimation highlighted the higher proportion of p24 compared with HIV-1 RNA in the 3K pellet (Figure 7D). This discrepancy was lower but still present in the 17K (Figure 7E) and 100K pellet (Figure 7F).

**Figure 7.**
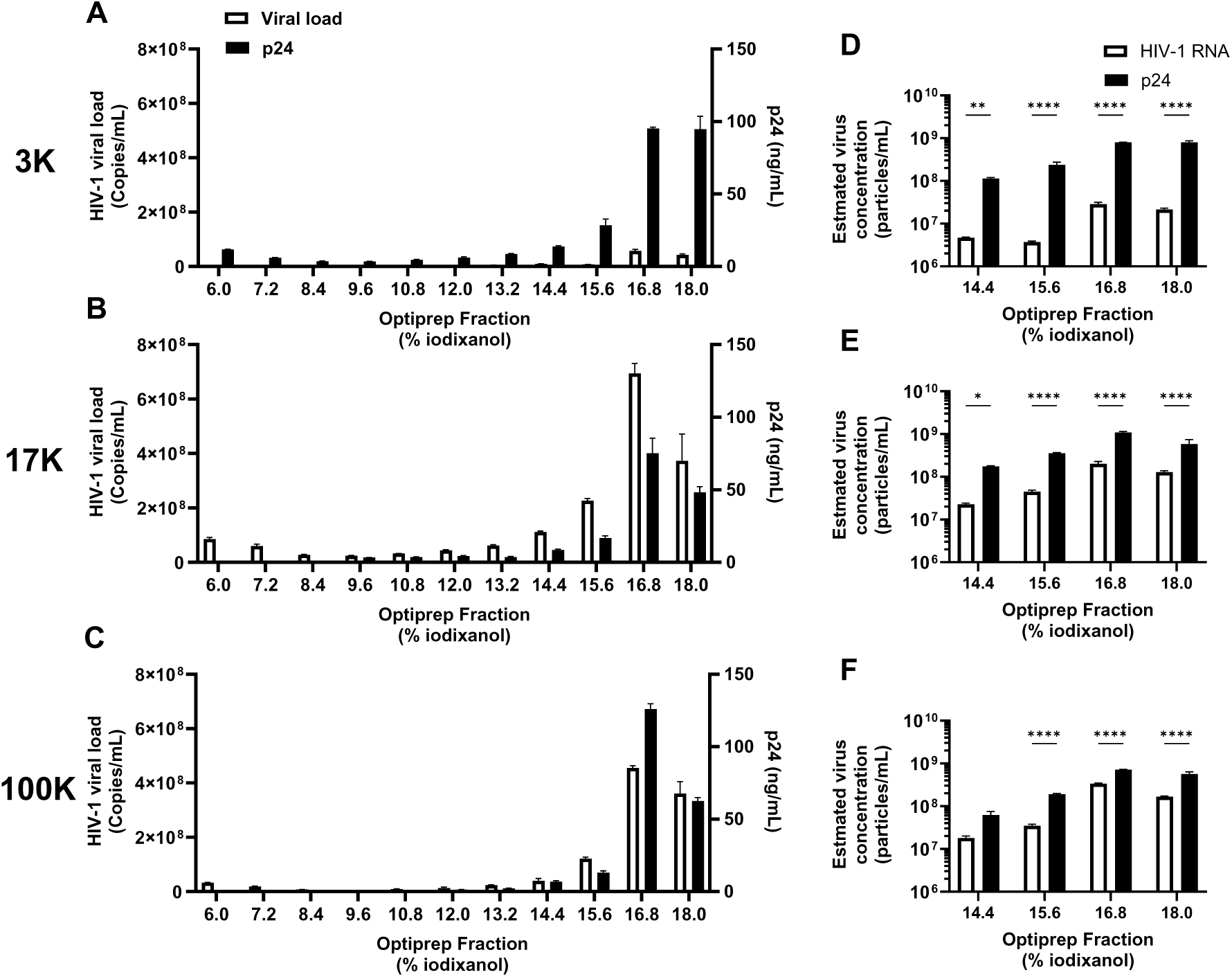
Viruses purification on an iodixanol velocity gradient. The 3K, 17K and 100K pellets from the supernatant of infected Raji CD4 DCIR cells were further purified on a velocity iodixanol gradient. The gradient comprised 11 fractions of iodixanol diluted in PBS ranging from 6.0 to 18.0% v/v with 1.2 % increments. Samples were deposited on the gradient and centrifuged at 180,000 x *g* for 50 minutes. HIV-1 RNA and capsid protein p24 concentrations were measured by RT-qPCR and ELISA in the 3K **(A)**, 17K **(B)** and 100K **(C)** pellets. Viral particles concentration estimated with the HIV-RNA and the p24 concentration in the 3K **(D)**, 17K **(E)** and 100K pellets **(F)**. Two-way ANOVA with Tukey’s multiple comparison test was performed for statistical analysis (* p < 0.05; ** p < 0.01; **** p < 0.0001).

## Discussion

The production of HIV-1 stock allows the study of HIV-1 infection in laboratory settings. HIV-1 stock contains EVs, and both particles are difficult to separate due to their biophysical similarities [9]. EVs’ effect on HIV-1 stock infectivity is generally disregarded, even if EVs can influence HIV-1 pathogenesis by transporting host and viral molecules [15, 16, 48, 49]. This study aimed to characterize the packaging of viral components in EV subtypes and decipher how the association of viral particles to each EV subtype affect their infectivity.

HIV-1 exits the infected cell by budding at the cell membrane or via multivesicular endosomes [10, 11]. EV content is controlled by its biogenesis pathway, the producing cell and the cell condition [50]. Our data showed viral RNA and capsid p24 protein packaging in EVs from the 3K, 17K and 100K pellets. Of note, p24 and HIV-1 RNA concentration did not correlate in the cell culture supernatant, which was unexpected because both molecules are produced during the virus’ replication cycle. The results with proteinase K-treated supernatants suggested that a significant proportion of HIV-1 RNA is not associated with p24 capsid production. The p24 protein was inside EVs, protected from proteinase K treatment. HIV-1 RNA concentration decreases upon proteinase K treatment, suggesting that a large proportion of EV-associated RNA is likely at the EVs’ surface. This might be caused by an incomplete replication cycle, leading to intracellular HIV-1 RNA accumulation [51], and HIV-1 RNA sorting in the extracellular environment. Extracellular HIV-1 RNA contributes to persistent immune activation and dysfunction through TLR sensing in dendritic cells, macrophages, and T cells [52, 53]. HIV-1 RNA on EV’s surface could participate in immune activation by the TLR pathway.

Estimations of viral particle concentrations were calculated with the HIV-1 RNA and p24 protein measurements. Our data showed an overproduction of p24 capsid protein compared to HIV-1 RNA, suggesting the presence of RNA-free viral particles. Simultaneous detection of genomic HIV-1 RNA and gp120 on a single-particle basis highlighted that 1 out of 8 viral particles contain genomic RNA [54], a similar ratio to our findings. These RNA-free capsids could contribute to the HIV-1 pathogenesis. Host restriction factors, such as tripartite-motif-containing 5α (TRIM5α), target the capsid to disrupt the virus replication cycle [55]. The abundant genomic RNA-free viral capsids might serve as decoys to TRIM5α, thus lowering the probability of TRIM5α binding to capsid-containing viral genomic material.

The most infectious virus produced by infected Raji CD4 DCIR pelleted at a lower speed with larger EVs such as apoptotic vesicles. Given that EVs from the 3K and 17K pellets were larger than HIV-1 particles (120-200 nm) [56], it is possible that one or many HIV-1 particles were within EVs. HIV-1 particles containing two or more capsids were previously reported [56]. Notably, multi-capsid particles were bigger than single-capsid viruses [56]. Whole and functional viruses have been reported to be within EVs. For example, enterovirus 71 is a non-enveloped virus that can exit infected cells non-lytically via the exosome biogenesis pathway [57]. EV-enclosed enterovirus 71 is protected from proteinase K digestion and maintains its infectivity [57]. The dengue virus can hide within EVs to evade antibody binding and recognition by the immune system [58]. Therefore, the integration of viral particles into larger EVs could be a mechanism utilized by HIV-1 to escape the immune system and facilitate its dissemination in tissues.

Another group found infectious particles associated with a large spectrum of EV fractions, even to extracellular particles smaller than exosomes. Their findings showed that viral particles associated with exomeres, particles smaller than exosomes, are the most infectious [21]. The second most infectious particles were in the larger EVs, apoptotic vesicles and microvesicles, while exosome-associated viruses were the least infectious [21]. Their results, obtained with a productively infected T-cell cell line (Jurkat), corroborate our observations with the myeloid/lymphoid hybrid Raji CD4 DCIR cell line. Conversely, HEK293T cell transfection produced more infectious viruses associated with smaller EVs. These results suggest that HEK293T transfection and Raji CD4 DCIR infection produce different HIV-1 particles, partly caused by their association with different EV subtypes. Moreover, Raji CD4 DCIR EVs and HIV-1 particles were positive for HLA-DR, ICAM-1 and miR-155, which HEK293T do not express [26–28]. HLA-DR, ICAM-1 and miR-155 enhance HIV-1 infectivity [23, 26, 34].

This study highlights the diverse range of viral particles and EVs released by infected Raji CD4 DCIR cells (Figure 8). The heterogeneous production of viruses and EVs in these cells may serve as a valuable model for more accurately replicating virus production in infected cells of PLWH. Further studies using primary cells are necessary to validate this hypothesis. A deeper understanding of the heterogeneity of viral particles generated during HIV-1 infection could improve virus production methods, enabling a more precise simulation of infection dynamics.

**Figure 8.**
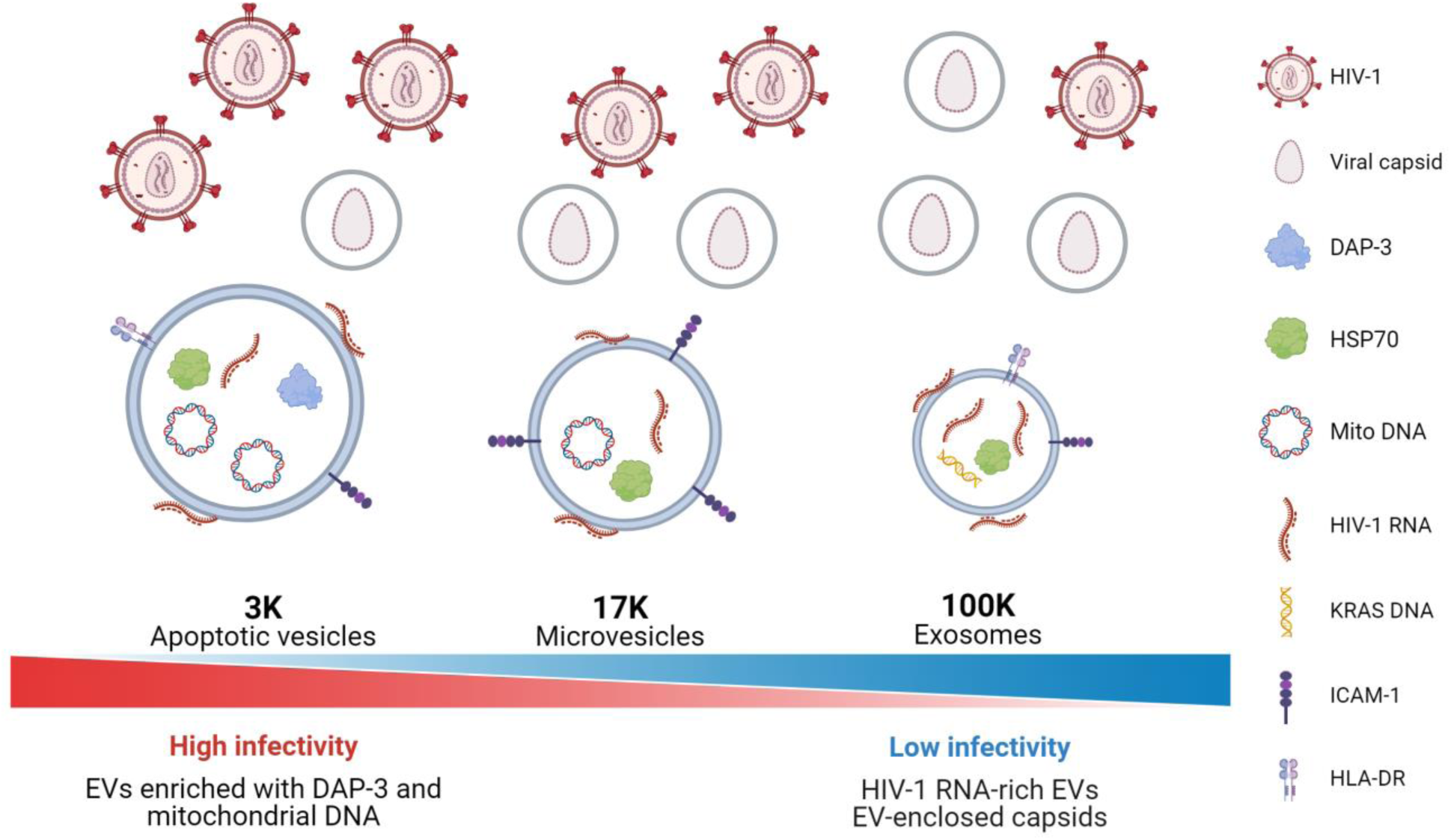
The diversity of EVs and viruses co-purified from infected Raji CD4 DCIR cells. The characterization of EVs and viruses co-purified by differential centrifugation. The 3K pellet contained viruses with high infectivity, along with DAP-3 and mitochondrial DNA enriched EVs. A lower infectivity was associated with the 17K pellet. The 17K EVs displayed ICAM-1 and contained HSP70 and mitochondrial DNA. The lowest infectivity was found in the 100K pellet, which likely contained HIV-1 RNA-rich EVs and genomic RNA-free viral capsids within EVs.

## Author contributions

Conceived and designed the experiments J.B. and C.G.: Performed the experiments: J.B. and A.R. Analyzed the data: J.B., A.R. and C.G.; Contributed clinical samples, reagents, materials, and analytical tools: J.B. and C.G.; Wrote the manuscript: J.B. and C.G. All authors have critically reviewed the paper and have agreed on the published version of the manuscript.

## Acknowledgements

This research was funded through Canadian Institutes of Health Research (CIHR) grants MOP-391232; MOP-188726; MOP-267056 (HIV/AIDS initiative) to C.G. The FRQ-S supports the Centre de recherche du CHU de Québec – Université Laval infrastructure. J.B. is the recipient of the Desjardins scholarship from the Fondation du CHU de Québec, a MITACS shoand the recruitment scholarship from the AIDS Research Fund of Université Laval. The authors thank Drs. Martin Pelletier and Stephane Gobeil for access to the qPCR platform and Dr. Éric Boilard for access to the nanoparticles flow cytometry platform. We thank Myriam Vaillancourt and Isabelle Allaeys from technical assistance. We thank Alexandre Bastien from the transmission electron microscopy observation at the plateforme d’imagerie de l’Université Laval. We thank Hend Jarras for proofreading the manuscript. We are very grateful to the study participants, without whom this study would not have been feasible.

## Institutional review board statement

The study was conducted according to the guidelines of the Declaration of Helsinki and approved by the Centre de Recherche du CHU de Québec-Université Laval (Québec, QC, Canada) ethics review boards C12-03-208 (2012-2021) and CER-2019-4258. All subjects were volunteers and provided written informed consent before participating in the study.

## Informed consent statement

Written informed consent was obtained from all subjects involved in the study.

## Data availability statement

The study protocol, results and informed consent documents will be made available to researchers upon request from the corresponding author. Researchers will be asked to complete a concept sheet for their proposed analyses to be reviewed, and the investigators will consider the overlap of the proposed project with active or planned analyses and the appropriateness of the study data for the proposed analysis.

