## Supplementaldata for "Characterization of HIV-1 particles co-purified with three extracellular vesicle subtypes from the Raji CD4 DCIR cell line, a hybrid model of CD4 T cells and dendritic cells"

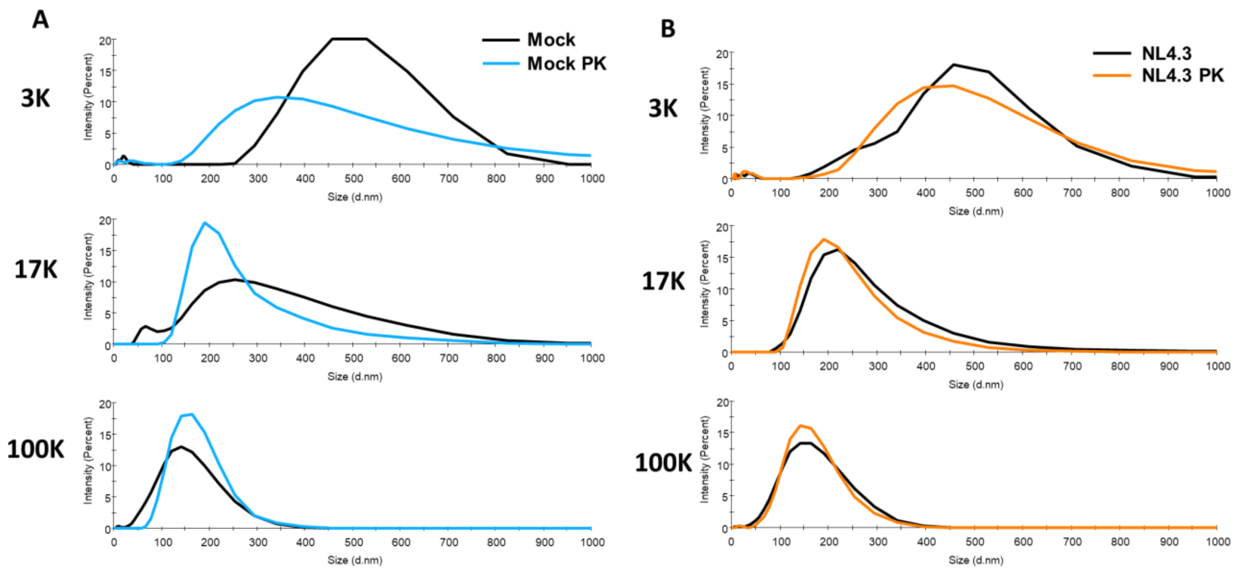


**Figure S1. Effect of proteinase K on particle size distribution.**
Hydrodynamic size was measured by DLS EVs from control cells (Mock) **(A)** EVs from infected cells (NL4.3) **(B)**. The presented data are the mean of the four EV/HIV-1 preparations.

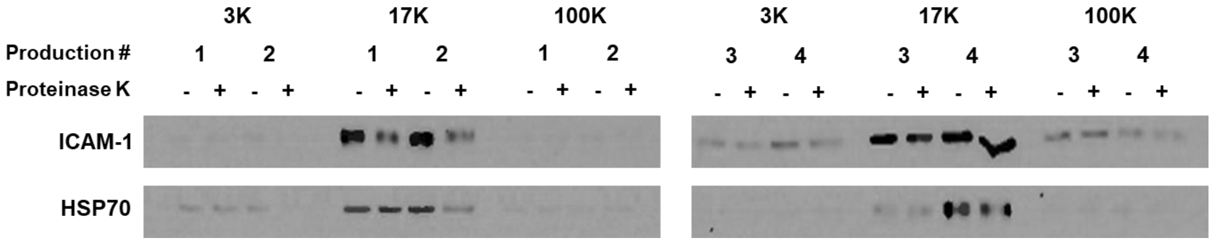


**Figure S2. Proteinase K effect on EV proteins.**
EVs protein markers from the 3K, 17K and 100K pellets of infected cells were characterized by Western Blot.

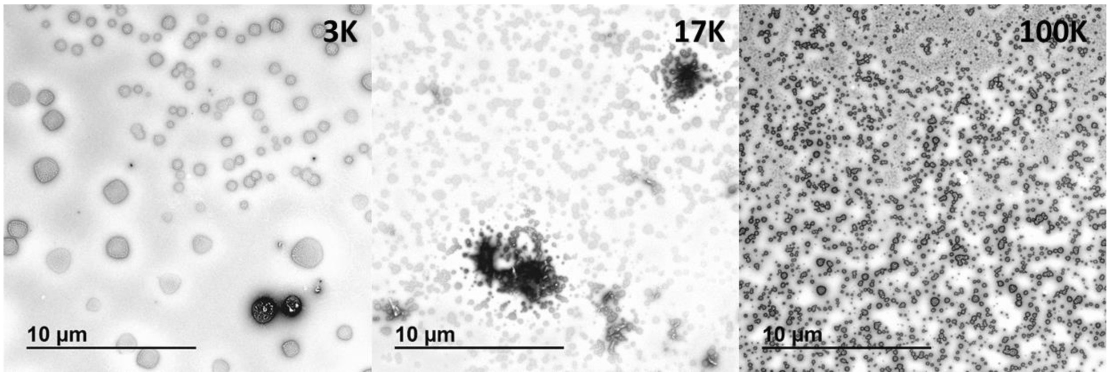


**Figure S3. Larger areas of transmission electron microscopy images.**
Transmission electron microscopy at a 1,000X magnification of pelleted EV/HIV-1 preparations.

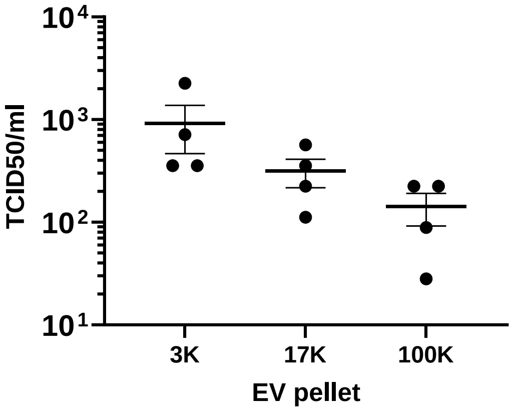


**Figure S4. Infectivity of HIV-1 particles on TZM-bl cells.**
HIV-1 infectivity in the 3K, 17K and 100K pellets was assessed with a TZM-bl indicator cell line. An equal volume of virus (45 µL) was used to infect the cells. Statistical analysis was carried out by one-way ANOVA.

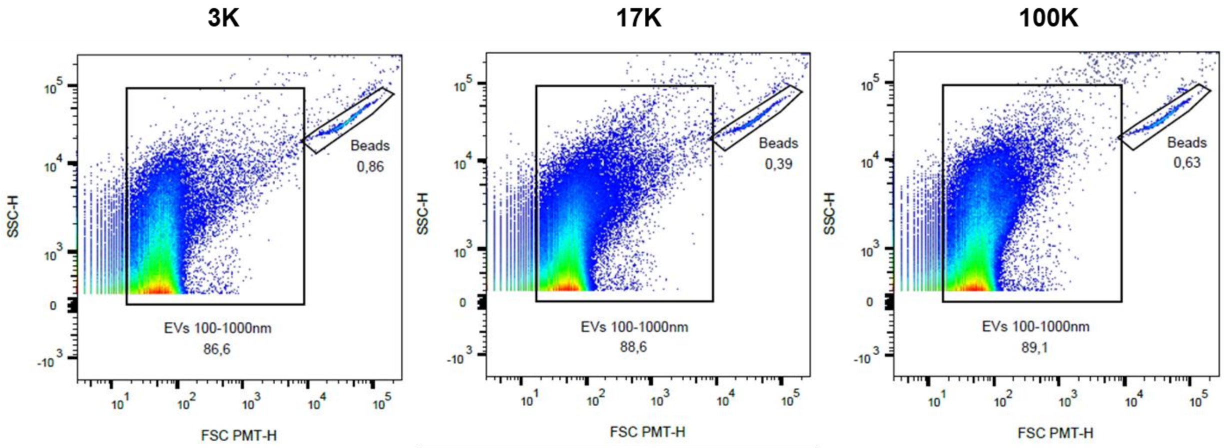


**Figure S5. Virus and EV size and complexity analysis by flow cytometry.**
A representative graphic of FSC-PMT (photomultiplicator) vs SSC of each EV and NL4.3 virus pellet from the supernatant of infected Raji CD4 DCIR cells.

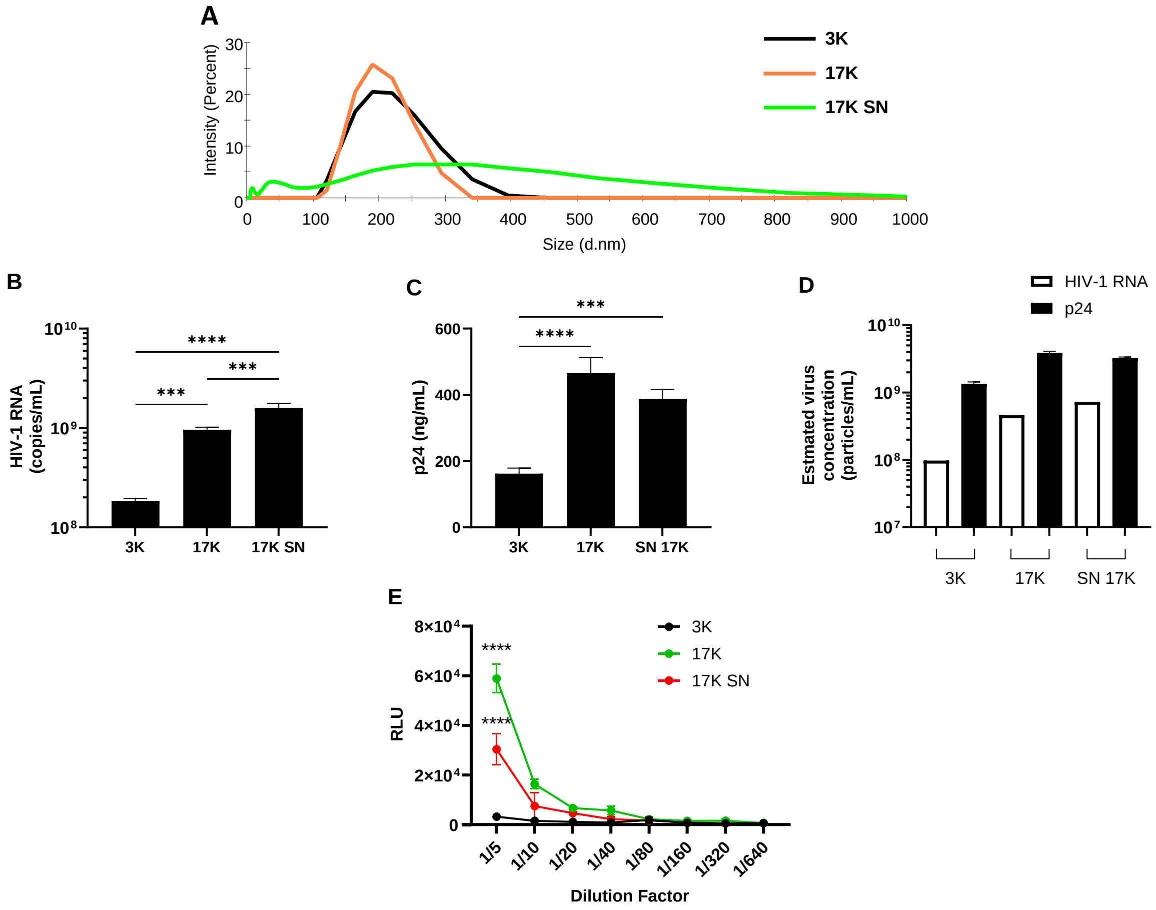


**Figure S6. Infectivity of HIV-1 particles associated with three EV fractions from transfected HEK293T cells.** EVs and virus hydrodynamic size measured by Nanosizer **(A)**. RT-qPCR quantification of viral RNA in the samples **(B)**. Capsid p24 protein quantification by ELISA **(C)**. Estimated virus concentration with the HIV-1 RNA and p24 concentrations **(D)**. Virus infectivity was measured with a TZM-bl cell line containing a luciferase gene controlled by HIV-1 transcription **(E)**. A two-way ANOVA was performed for statistical analysis (*** $p < 0.001$; **** $p < 0.0001$).

### Method description

#### Alternative purification of EV/HIV-1 produced by transfected HEK293T

EV/HIV-1 from the supernatant of transfected HEK293T were also purified using an alternative protocol. The supernatant was centrifuged at 3,000 x *g* for 15 minutes (3K pellet), and then at 17,000 x *g* for 30 minutes (17K pellet). Due to a technical limitation, the 100,000 x *g* (100K pellet) could not be performed for this protocol. Thus, the remaining supernatant after the 17,000 x *g* centrifugation (17K SN) was considered as the 100K pellet.

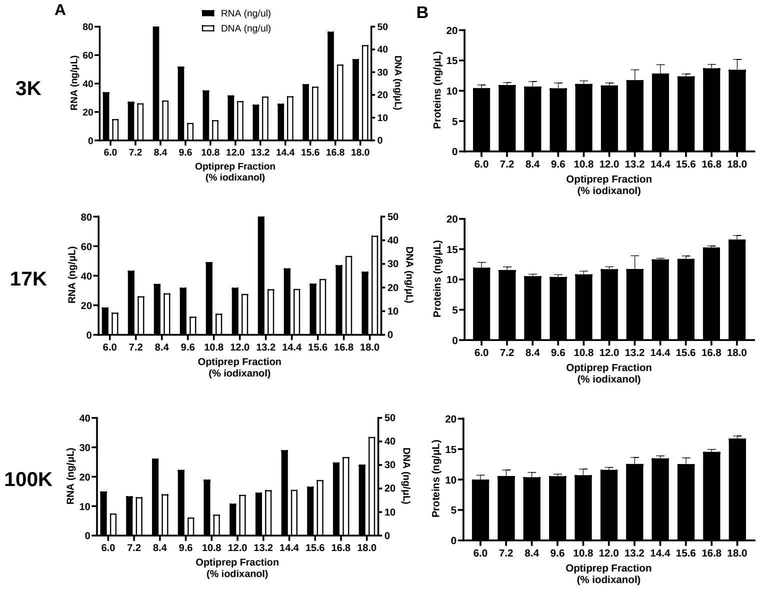


**Figure S7. Molecular content of each fraction from the iodixanol velocity gradient.**
The 3K, 17K and 100K pellets from the supernatant of infected Raji CD4 cells were further purified on a velocity iodixanol gradient. The gradient comprised 11 fractions of iodixanol diluted in PBS ranging from 6.0 to 18.0% v/v with 1.2 % increments. Samples were deposited on the gradient and centrifuged at 180,000 x *g* for 50 minutes. RNA and DNA distribution in the gradient **(A)**. Protein quantitation by microBCA assay **(B)**.

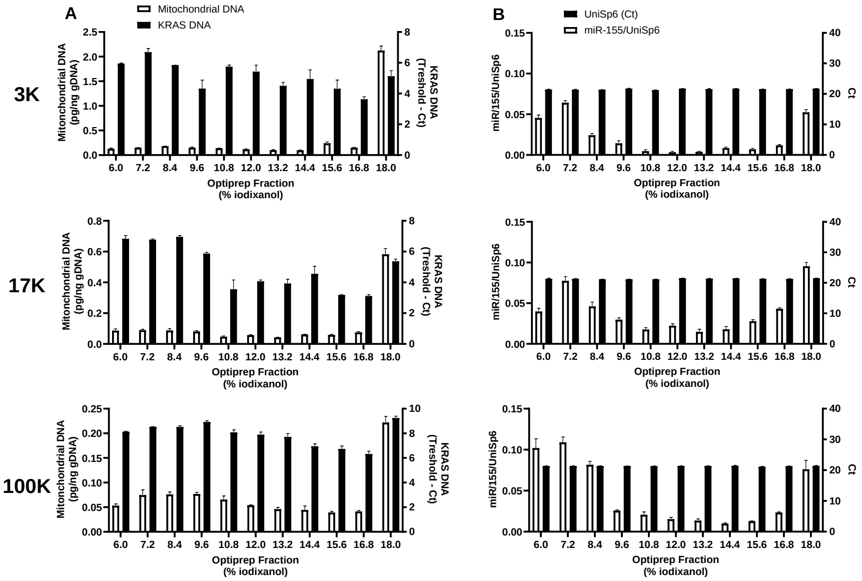


**Figure S8. Molecular content of each fraction from the iodixanol velocity gradient.**
The 3K, 17K and 100K pellets from the supernatant of infected Raji CD4 cells were further purified on a velocity iodixanol gradient. The gradient comprised 11 fractions of iodixanol diluted in PBS ranging from 6.0 to 18.0% v/v with 1.2 % increments. Samples were deposited on the gradient and centrifuged at 180,000 x *g* for 50 minutes. Mitochondrial and KRAS DNA quantification by qPCR. **(B)**. Normalization of miR-155 quantification in the gradient with the spike-in RNA UniSp6 **(B)**.
